# GSAn: an alternative to enrichment analysis for annotating gene sets

**DOI:** 10.1101/648444

**Authors:** Aaron Ayllon-Benitez, Romain Bourqui, Patricia Thébaut, Fleur Mougin

**Author notes:** These authors contributed equally (Patricia Thébaut, Fleur Mougin).

## Abstract

The revolution in new sequencing technologies, by strongly improving the production of omics data, is greatly leading to new understandings of the relations between genotype and phenotype. To interpret and analyze these massive data that are grouped according to a phenotype of interest, methods based on statistical enrichment became a standard in biology. However, these methods synthesize the biological information by *a priori* selecting the over-represented terms and may suffer from focusing on the most studied genes that represent a limited coverage of annotated genes within the gene set.

To address these limitations, we developed GSAn, a novel gene set annotation Web server that uses semantic similarity measures to reduce *a priori* Gene Ontology annotation terms. The originality of this new approach is to identify the best compromise between the number of retained annotation terms that has to be drastically reduced and the number of related genes that has to be as large as possible. Moreover, GSAn offers interactive visualization facilities dedicated to the multi-scale analysis of gene set annotations. GSAn is available at: https://gsan.labri.fr.

## 1 Introduction

Over the past decade, the revolution in new sequencing technologies has strongly supported the production of omics data to improve our understanding of the relations between genotype and phenotype. This research field involves analyzing gene sets to identify their biological function and to synthesize the key annotation information with the objective to help biologists in their interpretation. In this frame, many tools have been developed to support gene set analysis and visualization of annotations. Most of these tools are based on statistical enrichment methods that usually involve two stages: (i) an *a priori* stage that aims to synthesize the annotation by selecting the over-represented terms and (ii) an *a posteriori* stage to remove the potentially redundant information by using the Gene Ontology [1] relations. Examples of such enrichment-based tools are g:Profiler [2], clusterPro-filer [3] and WebGestalt [4]. In g:Profiler, statistically enriched terms are grouped if they share one or more common parent terms. Two filters, named *moderate* and *strong*, then make use of the hierarchical structure of the used ontologies. The specific functionality *simplify* in clusterProfiler provides a score to retain only the most statistically relevant enriched GO terms (obtained by the *EnrichGO* tool) according to semantic similarity measures. WebGestalt does not propose any step to reduce redundancy within enrichment results, but an annotation file free from redundancy may be used as input of the analysis. Other tools like DAVID [5] propose an *a posteriori* stage that clusters the annotation terms that may be related to each other according to the genes they annotate. This stage does not result in a reduction of terms but rather in a categorization of terms according to their use. The results are thus given as lists of related terms and an additional manual expertise is still required to synthesize the information. Moreover, significant limitations of enrichment-based methods have recently been reported [6, 7]. First, these methods tend to focus on the most studied genes and provide gene set annotation results that cover a limited number of annotated genes [8, 6, 7]. Moreover, visualization facilities often suffer from a lack of capacity to perform multi-scale analyses which may help users while interpreting their results.

To address these limitations, we developed a novel gene set annotation Web server, called *Gene set Annotation* (or GSAn). The implemented method uses semantic similarity measures that allow users to *a priori* reduce a large number of Gene Ontology terms by computing a synthetic annotation for a given gene set [9]. The originality of this new approach is to identify the best compromise between the number of retained annotation terms that has to be drastically reduced and the number of related genes that has to be as large as possible. Moreover, GSAn provides interactive visualization facilities dedicated to the multi-scale analysis of gene set annotations [10] and is available at: https://gsan.labri.fr.

## 2 GSAn method

GSAn is based on a method that annotates a gene set making use of the annotations from Gene Ontology Annotation (GOA) [11] and the hierarchical structure of Gene Ontology (GO) [1]. The method is composed of four steps (Figure 1).

**Figure 1:**
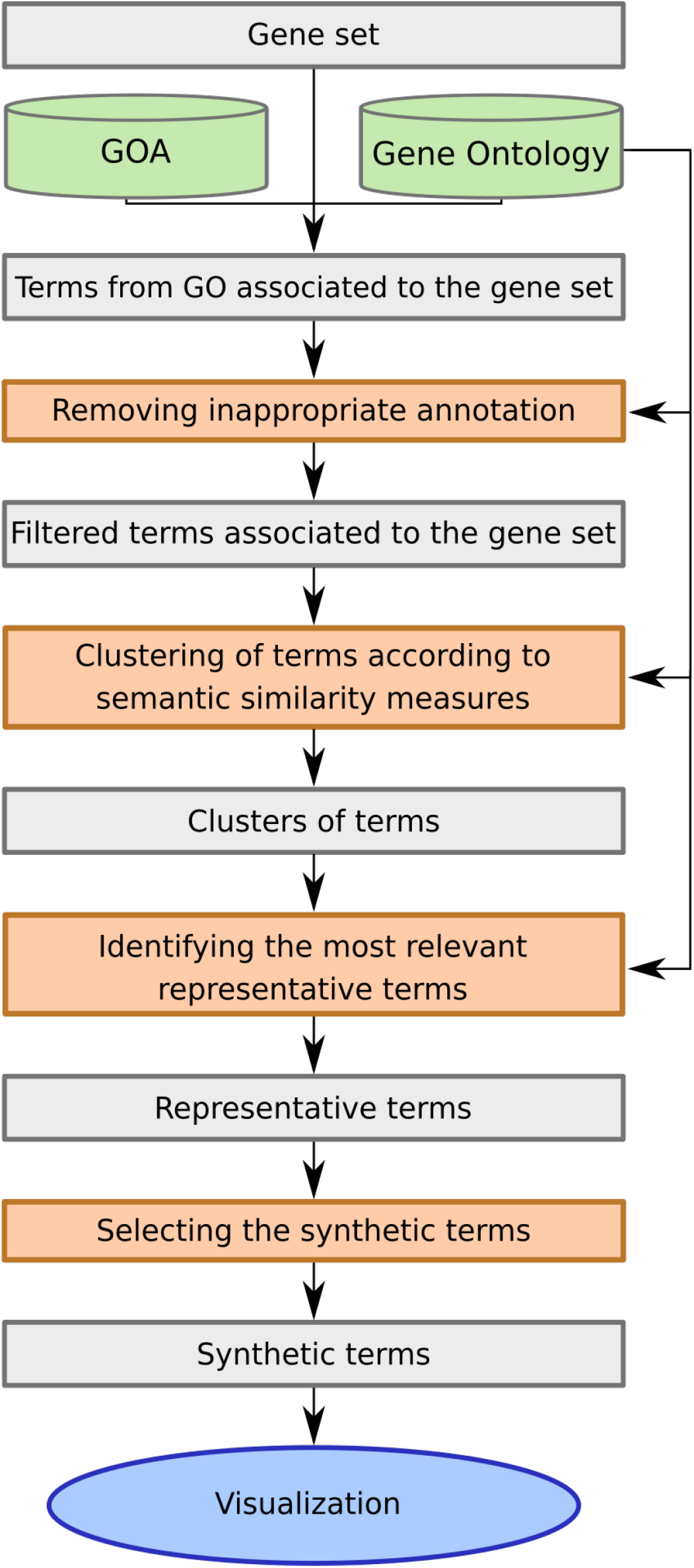
The GSAn method workflow. Steps are represented in orange rectangles while their input and output are displayed as grey rectangles.

### 2.1 Removing inappropriate annotation

First, we removed inappropriate annotations. An inappropriate annotation is defined by any association where the GO term does not provide relevant information. An annotation can be inappropriate for two reasons: redundancy and incompleteness. Criteria for removing redundant annotations are described in Supplementary data. The notion of incomplete annotation was first reported by Faria *et al.* [12] that considered terms with more than 10 descendants as inappropriate (being too general). We adapted this definition of incomplete annotation for taking into account the quality of the annotation. To this end, we considered the information content (IC) provided by Mazandu and Mulder [13] and computed the IC distribution of terms from GO. We then retained only GO terms whose IC was higher than the value of the first quartile.

### 2.2 Clustering of terms according to semantic similarity measures

The semantic similarity compares GO terms depending on ontological or annotation features. A pairwise semantic similarity measure is defined as a function that, given two terms, returns a value reflecting how close in meaning they are [14, 15]. A semantic similarity matrix was thus computed for each pair of GO terms associated with the gene set. The semantic similarity measures implemented in GSAn are: Resnik [16] normalized according to Jain and Bader’s approach [17], Lin [18], Aggregate Information Content (AIC) [19], NUnivers [13] and Distance Function [20]. Formulas of these semantic similarity measures are available in Supplementary Data. This matrix was then used to compute groups of terms according to the average linkage clustering algorithm (that exhibited the highest cophenetic correlation compared with other algorithms considered in [9]). The best number of clusters was determined using the Average Silhouette Width score [21].

### 2.3 Selecting the most relevant representative terms

We define a representative term as a term that exhaustively represents the various information given by the terms of a cluster. As the number of representative terms may vary according to the size of the cluster, two strategies were used to determine the best number. First, if a single term inside a cluster annotated more than 70% of genes, it was directly considered as representative. Secondly, if such a term did not exist, an algorithm described in [9] was applied to compute an appropriate number of representative terms for the cluster. At the end of this stage, a new set of terms is obtained from the addition of representative terms of each cluster.

Then, to retain the most relevant representative terms, we used two quality criteria: term redundancy and gene coverage.

#### Removing inappropriate representative terms

Some clusters of terms may have been generated from terms with low similarity between them, resulting in very general representative terms. We thus removed terms whose IC is lower than the first quartile. A new selection stage was then applied to eliminate potential redundancies. According to the type of hierarchical relationship (*is*_*a* or *part*_*of*), the removal of the ancestor terms may have a different impact on the number of annotated genes. To deal with this issue, a different strategy was applied according to the type of hierarchical relationships. For the *is*_*a* relationship, the representative terms being ancestors of other representative terms were removed. For the *part*_*of* relationship, only the parent or child terms annotating the largest number of genes were retained.

#### Filtering representative terms according to the gene coverage

To filter out the representative terms associated with a limited number of genes, we used a formula that depends on the size of the gene set used as input. The resulting filtering value gradually increases according to the number of genes (see the formula in Supplementary Data).

### 2.4 Selecting the synthetic terms

At last, a heuristic algorithm based on the set cover problem (SCP) [22] was applied to the representative terms for selecting the terms that best summarized the biological information within the gene set. Within this framework, we thus defined a solution of the SCP as a minimal set of terms covering the maximum number of genes of the gene set. Finally, to identify the terms considered as synthetic, a weight was assigned to each term for taking into account its IC and the number of genes it annotates. For a given term, this weight is defined as follows:

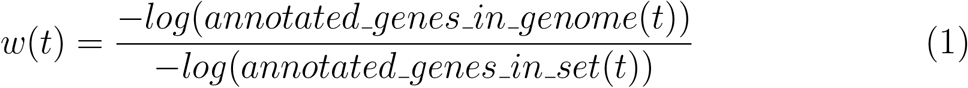

where *annotated*_*genes*_*in*_*genome*(*t*) and *annotated*_*genes*_*in*_*set*(*t*) correspond to the number of genes annotated by the term *t* within a whole genome and within a gene set, respectively. The numerator actually corresponds to the IC proposed by Resnik [16]. The pseudo-code of the synthetic algorithm (*SA*) and the customized implementation of SCP are described in Supplementary Data.

## 3 SERVER INTERFACE

### 3.1 Description input

At first, users have to upload a gene or gene product list and to select the appropriate organism within the form. Fourteen organisms are currently stored in GSAn, downloaded from the GO Web site^1^ and from the European Bioinformatics Institute Web site^2^, and listed in Supplementary Data. To be more flexible, users can also upload the annotation of any organism of interest using the *GAF* 2.1 format^3^. Users may choose any of the three GO sub-ontologies (being biological process or BP, molecular function or MF and cellular component or CC) or any combination of them. If more than one sub-ontology is chosen, the analyses are computed separately and results are then merged. Five semantic similarity measures are available (see the list in Methods). By default, GO annotations inferred automatically (evidence code: *IEA*) are included in the analysis but users may decide to exclude such annotations.

To customize the analysis, two advanced parameters are proposed to users: the gene support and the incomplete information filter. The gene support is the minimum number of genes that have to be associated to each representative term. The default value of this parameter is determined according to Formula (12) in Supplementary Data (based on the size of the gene set), and can be modified. The incomplete information filter is used to discard the terms presenting a low specificity in the ontology. Four levels of tolerance (none, low, medium and hard) can be applied corresponding to the percentile values given by the IC distribution (1, 10, 25 and 50 respectively) of GO terms. As a result, terms below the chosen percentile value are discarded. Optionally, users can provide their email address to be notified when the analysis is finished.

A summary of all of these parameters is given in Supplementary Data.

### 3.2 Description output

GSAn results are presented according to multiple visual metaphors. At the top left, three gauge plots display the global gene set information (see Figure 2A). The first one indicates the percentage of genes which are annotated by GO terms while the second one provides the percentage of genes part of GSAn results. Finally, the gene set similarity consists in a groupwise approach using the gene annotation proposed in [23]. A gene set similarity score of 1.0 means that all genes in the set have the same annotation and 0.0 means that terms have no common annotation. At the top right, a diverging bar plot display the gene coverage and the IC score of each synthetic term (see Figure 2B). Information about the representative terms is available within two separate pages in two different ways: a table (Figure 2C) and a combined tree visualization (Figure 2D). The table summarizes the information of each representative term, being synthetic or not. The tree visualization aims to describe the hierarchical context of each representative term within GO. To obtain such visualization, the GO structure (represented as a directed acyclic graph) was transformed in a tree according to the most informative parent of each representative term [10]. Two types of tree visualizations are then combined: a collapsible indented tree and a circular treemap. A tree color algorithm is applied to attribute similar colors to terms that are hierarchically related [24]. The brightness of the circle is related to the depth of terms in the ontology (darker means deeper in GO). White color forms represent the genes inside their annotation terms. Thus, a given gene can appear inside several terms of different branches. Moreover, within each gene circle, a bar chart is displayed to represent its annotation terms (using their assigned colors). This visualization allows to explore annotation results thanks to interactions such as zooming within the circular treemap, or expanding the branch in the indented tree (illustrated in Figure 2D). Additionally, users can download a JSON file and view these results again by uploading the file within the “Visualization” page. Also, results can be downloaded as a CSV format.

**Figure 2:**
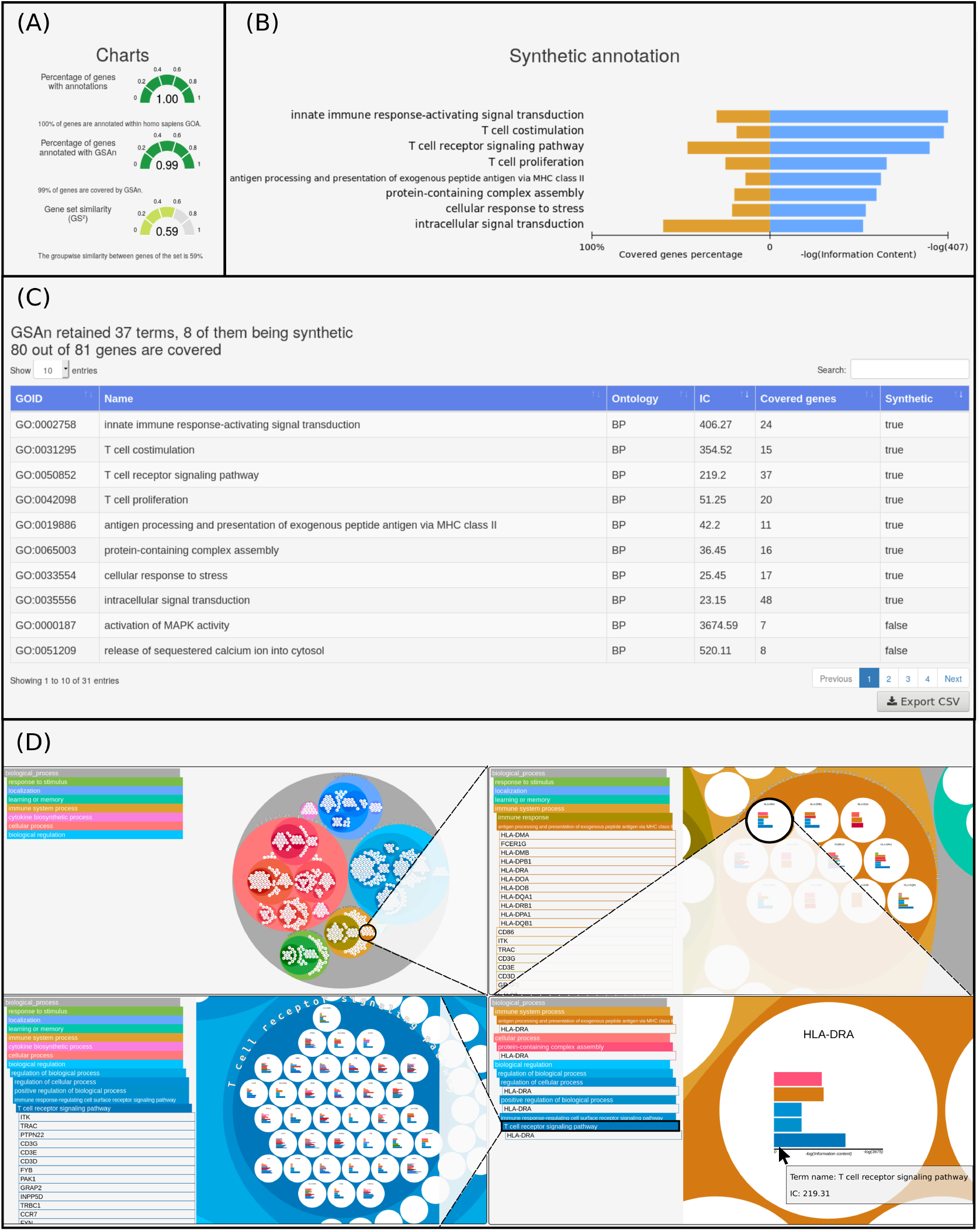
GSAn output results. (A) Three gauge plots show information about the annotated genes and the genes covered by GSAn as well as the groupwise similarity of genes in the set defined in [23]. (B) A diverging bar plot displays the IC and the gene coverage of each synthetic term. (C) A table presents all information about representative terms. (D) An example of the combined visualization shows 12 the click and zoom interactions.

### 3.3 Implementation and storing results

GSAn was implemented in JAVA EE using the SpringBoot framework. From the client side, the page exhibiting results was implemented based on JavaScript using the D3.js [25] and TreeColors.js^4^ libraries. The releases of GO and GOA are weekly updated. The JSON files created by GSAn are stored during 12 hours.

## 4 CASE STUDY

To illustrate GSAn, we present two analyses: (i) a comparison of the results with known enrichment tools (available in Supplementary Data) and (ii) an application of GSAn using a gene set involved in the immune system.

For both analyses, we used the dataset computed by Li *et al.* [26], called BTM for blood transcriptional modules. This dataset is a repertoire of 346 modules characterizing innate and adaptative immune response in vaccination studies and was built using a large-scale data integration of human blood transcriptome provided by the NCBI Gene Expression Omnibus. Moreover, “interactome”, “bibliome” and pathways extracted from public databases were integrated to create a set of transcription modules.

In the second case study, we use GSAn to analyze a BTM module annotated by experts as *regulation of antigen presentation and immune response* [26]. This module contains 81 genes involved in the signal transduction in the immunological process against pathogens. The default parameters of GSAn are used and the chosen semantic similarity measure is NUnivers. GSAn retain 37 representative terms covering 80 out of 81 genes and 8 of them are synthetic terms (Figure 2). The gauge plots show a high gene coverage using the GOA file (first gauge) and GSAn analysis (second gauge). At last, the third displays a gene set similarity of 0.59, which means that genes share a high number of terms.

By focusing on the synthetic annotation displayed within the diverging bar plot, we observe terms related to the proliferation and costimulation of T cell and the activation of signaling transduction by the innate immune response. Also, these terms and the term *antigen processing and presentation of exogenous peptide antigen via MHC class II* (GO:0019886) are consistent with the manual annotation performed by experts and show that the annotation provided by GSAn is even more specific. Indeed, GSAn illustrates that the module is also involved in the proliferation of T cell. Moreover, more complete information may be observed from the representative terms through the information table or the combined tree visualization. By exploring the tree visualization, we obtain additional information, such as terms sharing the same informative ancestor or the genes annotated by more than one term. For example, by focusing on the term *antigen processing and presentation of exogenous peptide antigen via MHC class II*, we notice that eleven genes are annotated by this term. When clicking and developing in details each gene, we observe that six out of the eleven genes are annotated by *T cell receptor signaling pathway* (GO:0050852) and three of them by *T cell proliferation* (GO:0042098). Thus, with very few user interactions, we retrieve additional information about the biological role of some genes in the module.

## 5 CONCLUSION

The main problems in finding gene signatures are mainly related to the investigation of the biological function of gene sets. That problem can be solved using classical enrichment methods, such as DAVID or g:Profiler. However, these methods focus on the most studied genes that may provide annotations covering a limited number of annotated genes [8, 6, 7]. Another problem is the redundant information within annotations that may increase the difficulty in interpreting results when no *a posteriori* analysis is performed. To address these issues, we propose a new Web server as an alternative to classical enrichment analysis. The underlying method makes use of the hierarchical structure of GO to reduce the number of terms while keeping an appropriate level of biological information. Compared to enrichment analysis tools, GSAn has shown excellent results in terms of maximizing the gene coverage while minimizing the number of terms. GSAn has provided a gene set annotation that is more specific than results given by experts (for a human gene set). Also, an originality of GSAn is to provide interactive visualization abilities to analyze the resulting gene set annotations. The underlying visualization is based on a combined tree that supplies zoom operations to browse terms and the genes they annotate according to the level of biological information that may interest users.

## Footnotes

1 http://www.geneontology.org/page/downloads

2 https://www.ebi.ac.uk/GOA/downloads

3 http://www.geneontology.org/page/go-annotation-file-gaf-format-21

4 https://github.com/e-/TreeColors.js/

