## Supplementary notes for "GSAn: an alternative to enrichment analysis for annotating gene sets"

### Supplementary data

Aarón Ayllón-Benítez, Patricia Thébault, Romain Bourqui, Fleur Mougin

**3 Comparing GSAn to enrichment tools** **10**

### 1 Supplementary details of the GSAn method

#### 1.1 Removing inappropriate annotation

In addition to remove incomplete annotation, we also eliminated redundant annotation.

Two features can help to identify redundancy: associated evidence and term relationships. For the first feature, GOA provides an evidence code for each gene-term association that explains how this annotation was acquired. Additional information such as *NOT*, *contributes_to* and *colocalizes_with* are specified for qualifying some annotations. Thus, the annotations that were exclusively associated with the evidence code *ND* (no biological data available) or with the qualifier *NOT* were removed. As for the second feature, redundancy corresponds to situations where a gene is annotated by two GO terms hierarchically related. In such case, only the association involving the most specific GO term was retained. Moreover, considering the regulatory relationships within GO, we assumed that a gene associated with a term that regulates another term is also involved in the regulated term. Thus, all regulation terms have been replaced by their regulated terms. For example, the term *regulation of ion transport* (`GO:0043269`) was replaced by *ion transport* (`GO:0006811`).

#### 1.2 Pseudocode to identify the synthetic terms

During the last step of GSAn (see the subsection `Selecting the synthetic terms` from the GSAN METHOD section of the main paper), the synthetic terms have to be identified. Thus, to recover them while preserving the same number of covered genes, we applied the following heuristic algorithm that is based on the set cover problem (SCP).

**Algorithm 1:** $SA(R, G)$

**Input** : $R$ is a set of representative terms,

$G$ is a set of genes covered by at least one term from $R$.

**Used functions**: $genes(x)$ is the list of genes in $G$ annotated by the term $x$,

$w(x)$ is the weight score of the term $x$, determined as follows:

$$w(x) := \frac{-log(annotated\_genes\_in\_genome(x))}{-log(annotated\_genes\_in\_set(x))}$$

1) Let $S$ represent the set of synthetic terms and $C$ represent the set of genes covered by all terms in $S$.

$Initialize\ S := \emptyset$

$Initialize\ C := \emptyset$

2) While $C$ is not the same as $G$ :

$a$) Find the term $r \in R$ whose score is the biggest, in which score is defined by:

$score(r) := |genes(r) - C| \cdot w(r)$

$b$) Add the term with the biggest score to $S$ and remove it from $R$

$S := S \cup \{r\}$

$R := R - \{r\}$

$C := \forall s \in S| \bigcup genes(s)$

**return** $S$

### 2 GSAn input parameters

The input parameters to be used in GSAn for the analysis are listed in Table 1. The next subsections provide details about the organisms and the semantic similarity measures available in GSAn as well as the way the gene support is calculated.

Table 1: Input parameters to be used in GSAn for the analysis

| Parameter | Description | Default value |
|---|---|---|
| Gene list | A list of gene identifiers | - |
| Genome annotation | Organism name that will be used to recover the gene annotations. Fourteen organisms are proposed and any other organism can be uploaded using the GAF 2.1 format. | homo_sapiens |
| Inferred from electronic annotation (IEA) | Boolean that indicates whether the IEA annotations should be used | true |
| Ontology | Sub-ontology of GO (BP, MF, CC) to be used for the analysis | BP |
| Semantic similarity measure | Measure to be used for computing the term similarity matrix | AIC |
| **Advanced parameter** | | |
| Gene support | Minimal number of genes that have to be covered by any representative term | see Formula 12 |
| Incomplete information filter | Tolerance degree to discard terms with a low specificity | medium |
| Email | Provided by users for being notified when the analysis is finished. | - |

#### 2.1 List of organisms available in GSAn

| Organism | File |
|---|---|
| *Sus scrofa* | goa_pig.gaf |
| *Saccharomyces cerevisiae* | goa_yeast.gaf |
| *Rattus norvegicus* | goa_rat.gaf |
| *Mus musculus* | goa_mouse.gaf |
| *Homo sapiens* | goa_human.gaf |
| *Gallus gallus* | goa_chicken.gaf |
| *Escherichia coli* | gene_association.ecocyc |
| *Drosophila melanogaster* | goa_fly.gaf |
| *Danio rerio* | goa_zebrafish.gaf |
| *Canis lupus* | goa_dog.gaf |
| *Candida albicans* | gene_association.cgd |
| *Caenorhabditis elegans* | goa_worm.gaf |
| *Bos taurus* | goa_cow.gaf |
| *Arabidopsis thaliana* | goa_arabidopsis.gaf |

#### 2.2 Semantic similarity measures implemented in GSAn

Before presenting the semantic similarity measures available in GSAn, we first introduce the notion of information content because it is used in most of these measures.

##### 2.2.1 Information content

The information content, or IC, is a score associated with a term within an ontology, indicating how much this term is informative. The bigger the IC is, the more specific the term is. Two kinds of IC are used in GSAn: *intrinsic* and *extrinsic*. The intrinsic IC uses only information available within the ontology structure and the extrinsic IC uses external information.

**Extrinsic IC.** The extrinsic IC that is used in GSAn was proposed by Resnik in 1995 [1]. The principle is to compute the frequency of occurrence of a word in a given document. Applied to the Gene Ontology (GO), the frequency of a GO term is determined by the number of genes that are annotated by a GO term within the GOA file of a given organism. Thus, the probability of a GO term is computed as follows:

$$p(t) = \frac{frequency(t)}{frequency(root)} \tag{1}$$

Thus, the extrinsic IC is computed as follows:

$$IC_{ext}(t) = -log(p(t)) \tag{2}$$

This IC is used by some semantic similarity measures, whose formulas are provided in the next subsection.

**Intrinsic IC.** The intrinsic IC used in GSAn is the one proposed by Mazandu and Mulder that takes into account the position of GO terms in the GO structure [2]. First, the probability is computed recursively, from the root term to the leaf terms.

Each term depends on the probability of its parents divided by the children of the parent. Then, the IC provided by Mazandu for a term $t$ is the following:

$$p(t) = \begin{cases} 1 & \text{if } t \text{ is a } root. \\ \prod_{t_p \in parents(t)} \frac{p(t_p)}{|children(t_p)|} & \text{otherwise.} \end{cases} \quad (3)$$

$$IC_{GOu}(t) = -log(p(t)) \quad (4)$$

**Hybrid IC.** Song *et al.* [3] proposed a hybrid IC combining the semantic value of Wang *et al.* [4] and the extrinsic IC of Resnik [1]. The semantic value and the semantic weight are computed as follows:

$$SW(t) = \frac{1}{1 + e^{-\frac{1}{IC_{ext}(t)}}} \quad (5)$$

$$SV(t) = \sum_{t \in ancestors(t)} SW(t) \quad (6)$$

##### 2.2.2 Semantic similarity measures

**Resnik.** This semantic similarity measure was proposed by Resnik simultaneously with its IC [1]. The similarity between two terms corresponds to the IC of their most informative common ancestors (currently designated as $MICA$). The equation, after being normalized according to Jain and Bader's approach [5], is as follows:

$$Sim_{Resnik}(t_a, t_b) = \max_{t_{anc} \in ancestors(t_a) \cap ancestors(t_b)} (IC_{ext}(t_{anc})) \quad (7)$$

where $ancestors(t_x)$ and $IC_{ext}(t_x)$ correspond respectively to the ancestor terms and the extrinsic IC score of the term $t_x$.

**Lin.** To improve the results of Resnik's similarity, Lin proposed the following normalization [6].

$$Sim_{Lin}(t_a, t_b) = \frac{2 \cdot Sim_{Resnik}(t_a, t_b)}{IC_{ext}(t_a) + IC_{ext}(t_b)} \tag{8}$$

**Aggregate IC (AIC).** The aggregate information content or AIC is an alternative measure proposed by Song *et al.* using the hybrid IC [3]. The semantic similarity is thus computed according to the similarity provided by Wang *et al.* [4] as follows:

$$Sim_{AIC}(t_a, t_b) = \frac{\sum\limits_{t_{anc} \in ancestors(t_a) \cap ancestors(t_b)} 2 \cdot SW(t_{anc})}{SV(t_a) + SV(t_b)} \tag{9}$$

**NUnivers.** Using the IC proposed by Mazandu and Mulder [2], the similarity of NUnivers between two terms is the IC of the most informative content ancestor divided by the maximal IC between the compared terms.

$$Sim_{NUnivers}(t_a, t_b) = \frac{\max\limits_{t_{anc} \in ancestors(t_a) \cap ancestors(t_b)} IC_{GOu}(t_{anc})}{\max\{IC_{GOu}(t_a), IC_{GOu}(t_b)\}} \tag{10}$$

**Distance Function.** The distance function (DF) is adapted from the Jaccard index [7], as follows:

$$Sim_{DF}(t_a, t_b) = \frac{|ancestors(t_a) \cap ancestors(t_b)|}{|ancestors(t_a) \cup ancestors(t_b)|} \tag{11}$$

#### 2.3 Formula to determine the gene support

As indicated in the subsection `Identifying the most relevant representative terms` from the GSAN METHOD section of the main paper, we filter out the representative terms associated with a few number of genes. This corresponds to the gene support that an advanced input parameters of GSAn. Thus, we use a formula that depends on the size of the gene set used as input. The resulting filtering value gradually increases according to the number of genes. This formula is the following:

$$f(gs) = \begin{cases} 2 & \text{if } |gs| < 10. \\ floor(\sqrt{|\frac{|gs|}{10} - 1|}) + 2 & \text{otherwise.} \end{cases} \quad (12)$$

where $gs$ is a gene set. This filtering value computes the minimal number of genes for a given gene set.

### 3 Comparing GSAn to enrichment tools

GSAn has been compared to the classical enrichment analysis tools: g:Profiler [8], clusterProfiler [9], WebGestalt [10] and DAVID [11] (Table 2). As mentioned in the introduction, each tool provides a reduction stage to reduce the number of terms by eliminating the redundancy, except for DAVID. This comparison investigates the impact of the reduction step to decreasing the number of annotation terms while maintaining the number of annotated genes.

Table 2: Comparison of gene set functional analysis tools.

| | GSAn | DAVID | g:Profiler | clusterProfiler | WebGestalt |
|---|---|---|---|---|---|
| Statistical method | - | Fisher's Exact | Hypergeometric | Hypergeometric | Hypergeometric |
| Genes id | Symbol | Many | Many | Many | Many |
| Accessibility | WebSite, Rest Api | WebSite, Rest Api, R | WebSite, Rest Api, R | R | WebSite, R |
| Organism number | 14* | hundreds | hundreds | 2 | 12* |
| Updates | daily | - | - | - | - |
| Other resources | No | Yes | Yes | Yes | Yes |
| Reduction step | Synthetic algorithm, see Method for more details | - | Hierarchical filtering of the terms based on the p-value | Score filtering of the terms based on semantic similarity measures | Hierarchical filtering of the terms in the annotation file before computing the analysis |

* The addition of any organism annotation file is possible.

To carry out this comparative analysis, we focused on the GO biological process terms and retained only the BTM modules [12] for which a gene set annotation was provided by all tools. 226 modules were thus considered for the analysis. An IC distribution measure was computed based on the IC proposed in [2] and the ontology. This measure is relevant to analyze the gene coverage and the number of terms providing by each tool. We used four thresholds (from $Q_0$ to $Q_3$) corresponding to the quartiles of the IC distribution to filter the results of each tool. Each threshold filters out the terms with an IC below their value. Thus,

$Q_0$ refers to the whole resulting GO terms and $Q_1$, $Q_2$ and $Q_3$ correspond to GO terms having an IC value over 18.4, 44.4 and 155.3, respectively.

Figure 1 displays, for each tool, the gene coverage and the number of terms according to the IC distribution measure.

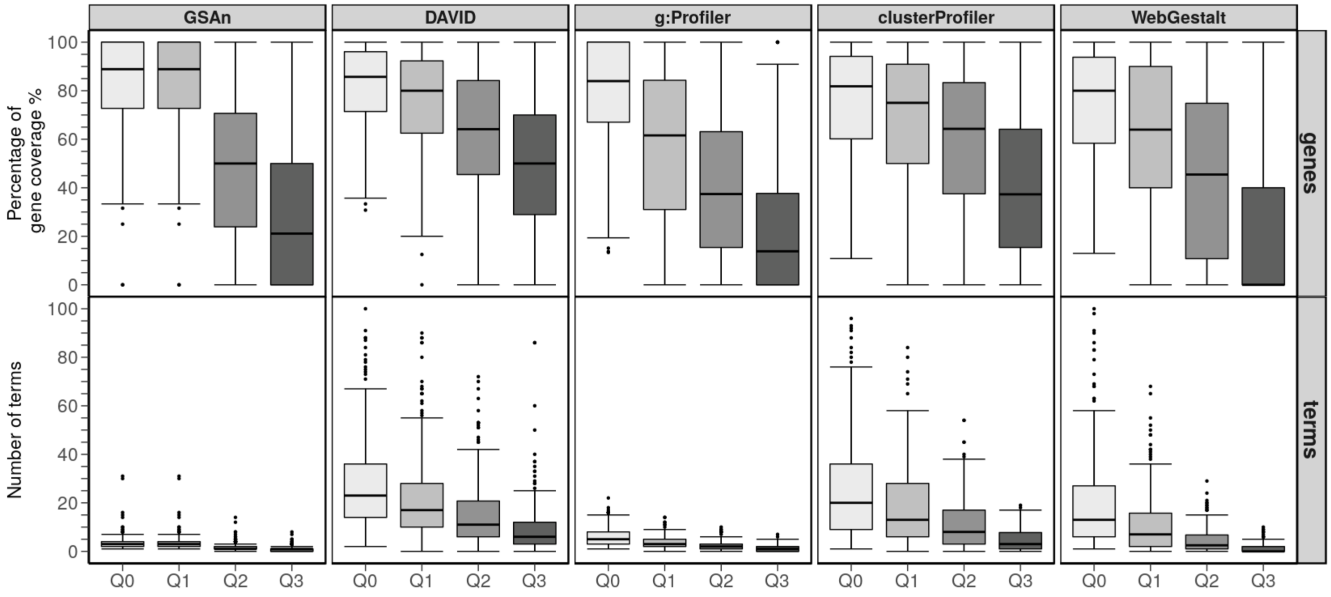

Figure 1: Box-plots providing the impact of the gene coverage percentage and the number of terms for each tool using the BTM modules [12] according to the quartile computed by the IC distribution in GO. Each quartile $Q_x$ corresponds to the IC value according to which the terms are filtered out. Thus, $Q_0$ corresponds to the whole set of terms provided by the tools and $Q_3$ to terms with an IC value higher than the third quartile of the IC distribution in GO.

Regarding the distribution of the gene coverage in $Q_0$, we do not observe any significant difference between tools. Thus, all tools present a median value higher than 80% of the complete gene set. On the other hand, regarding the number of terms, two classes of tools can be identified according to $Q_0$. First, DAVID,

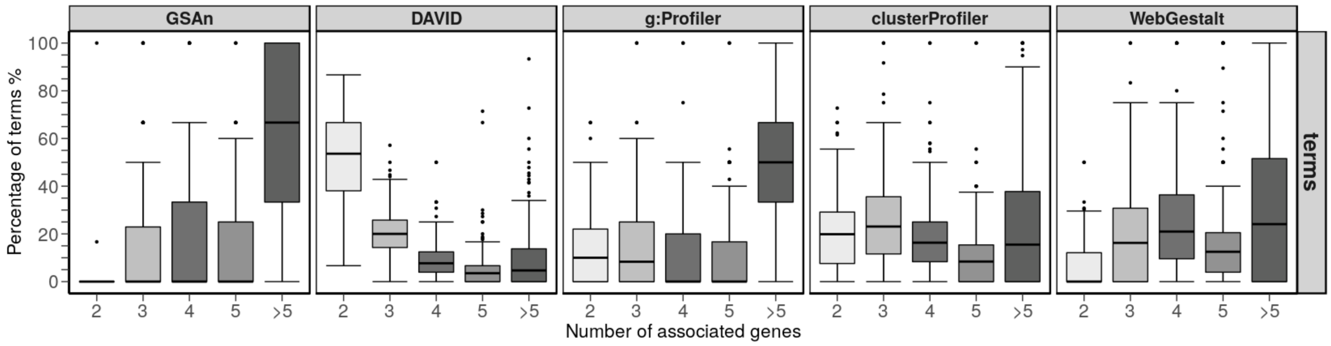

Figure 2: Box-plots showing the classification of terms by 2, 3, 4, 5 and more than 5 annotated genes. When terms are annotated mainly by 2 or 3 terms, it means that the result presents more specific terms. When terms are annotated mainly by more than 5 terms, it means that the terms are more general but that they are involving more genes.

clusterProfiler and WebGestalt give a median number of terms ranging from 10 to 25. The second group, involving GSAn and g:Profiler, has a smaller dispersion of numbers according to the various gene set with a median value ranging from 0 to 5. This smaller number of terms combined with a high gene coverage is relevant because very few terms annotate almost the whole genes. Considering $Q_1$, all tools except GSAn get decreased gene coverage and number of terms. GSAn keeps the same values as for $Q_0$ due to the IC filter applied to remove the incomplete information (corresponding to the first quartile in the distribution).

At last, $Q_2$ and $Q_3$ present the results for the most specific terms. The gene coverage is higher for DAVID and clusterProfiler with a median of gene coverage over 40%. They both lead to a better compromise regarding the number of terms and the gene coverage while maintaining relevant knowledge. However, the high number of terms may suggest that each resulting term is likely to annotate few genes.

To further investigate this hypothesis, we analyze the percentage of terms according to the number of genes that are annotated by these terms. Thus, Figure 2 shows the percentage of terms annotating 2, 3, 4, 5, and more than 5 genes. We observe that a median value of 50% of terms provided by DAVID annotate only two genes and the rest of the median boxes do not exceed 20%. On the contrary, the majority of terms provided by GSAn and g:Profiler annotate more than five genes. That suggests the existence of some general terms among GSAn and g:Profiler results that involve more genes compared to DAVID and clusterProfiler that include terms annotating few genes. Lastly, clusterProfiler and WebGestalt follow a similar behavior, with a median value for each box under 25%.
